# Oxytocin enhances acquisition in a social trust task in mice, whereas both oxytocin and its antagonist block trust violation learning

**DOI:** 10.1101/2024.12.13.628349

**Authors:** Samuel Budniok, Zsuzsanna Callaerts-Vegh, Marian Bakermans-Kranenburg, Guy Bosmans, Rudi D’Hooge

**Affiliations:** Laboratory of Biological Psychology, University of Leuven, Tiensestraat 102, Leuven, Belgium; Learn2Trust Research Group, University of Leuven, Tiensestraat 102, Leuven, Belgium; William James Center for Research, ISPA - University Institute of Psychological, Social and Life Sciences, Lisbon, Portugal; Facultad de Psicología y Humanidades, Universidad San Sebastián, Chile

## Abstract

The complex effects of the neurohormone oxytocin (OT) on socio-cognitive phenomena have recently been proposed to be complementary with safety learning, where a stimulus acquires safety-predicting properties when it predicts non-occurrence of an aversive event. OT may enhance saliency of safety stimuli and promote positive social behavior, such as trust, by reducing anxiety and stress. Complementary, OT may reduce the ability to modulate previously learned behaviors based on new, contradicting information. This occurs through its attenuation of prediction error (PE)—the discrepancy between expectations and actual outcomes. In the current study, we stimulated or inhibited (with antagonist cligosiban, CL) the OT system and subjected male and female mice to our social transmission of food preference (STFP) protocol to assess social safety learning. STFP is based on the observation that food neophobia of rodents is attenuated when a conspecific signals the safety of the food. We used safe food preference as putative murine homologue of human trust acquisition, and modeled trust violation (PE) using lithium chloride (LiCl)-induced food aversion after social interaction. In males, results revealed that OT enhanced trust acquisition, whereas both OT and its antagonist CL similarly blocked trust violation learning. None of the manipulations affected female behavior. Our findings highlight the complexities of OT’s role in social behavior, emphasizing caution in therapeutic manipulations of this system.

## Introduction

The effects of the neurohormone oxytocin (OT) on socio-cognitive phenomena turn out to be more complex than initially thought. The OT peptide is synthesized in paraventricular (PVN) and supraoptic (SON) nuclei of the hypothalamus, and released in the bloodstream to regulate birth-related processes such as lactation and labor (Shyken & Petrie, 1995; Uvnäs-Moberg et al., 2020). Additionally, neuromodulatory OT is released from PVN and SON terminals in various brain regions to regulate psychosocial concomitants of parturition, such as maternal bonding (Dulac et al., 2014), as well as broader aspects of socio-cognitive functioning, such as attachment, social recognition and aggression (Beery, 2015; Heinrichs & Domes, 2008). The involvement in these diverse social behaviors has recently been proposed to be complementary with safety learning (Eckstein et al., 2019). During safety learning, organisms learn that a neutral stimulus acquires safety-predicting properties when it predicts non-occurrence of an aversive event (Laing et al., 2024). OT is thought to modulate perceived safety, depending on context and individual learning history (Eckstein et al., 2019). Different mechanisms have been proposed to explain this effect.

Firstly, OT may enhance saliency of safety stimuli and promote positive social behavior by reducing anxiety and stress (Eckstein et al., 2019; Shamay-Tsoory & Abu-Akel, 2016). For instance, Smith and Wang (2014) demonstrated in prairie voles that social support reduced biobehavioral responses to stress, which could be mimicked by OT infusion in PVN, and abolished by an OT antagonist. Also, OT microinfusion into medial prefrontal cortex (mPFC) of rats enhanced fear extinction, whereas an antagonist impaired extinction (Brill-Maoz & Maroun, 2016). Finally, OT release during social support in rats recruits central amygdala neurons to switch fear stimuli to safety stimuli (Hegoburu et al., 2024). In turn, OT’s modulation of fear and stress may facilitate positive behaviors in safe contexts, such as maternal and pair bonding (Bosch & Neumann, 2012; Loth & Donaldson, 2021).These findings converge with the effect of OT on social support-seeking in humans, OT release during support, and its contribution to human trust and attachment (Bakermans-Kranenburg & Van IJzendoorn, 2013; Cardoso et al., 2013; Cuyvers et al., 2024; Shorey et al., 2023).

Complementary to saliency coding, some authors explained the effects of OT by its influence on prediction error (PE) processing (Ide et al., 2018). This refers to the difference between expectations about an event and its actual outcome that drives learning (Diederen & Fletcher, 2021). Intranasal OT promotes both unjustified trust and distrust in humans by attenuating PE processing (Ide et al., 2018). In other words, when participants initially learned to trust or distrust, OT discounted the weight of subsequent contradictory learning experiences.

In the current study, we stimulated or inhibited (with antagonist cligosiban, CL) the OT system during social safety learning in mice (as putative murine homologue of human trust acquisition). In line with the proposed mechanisms (Ide et al., 2018; Shamay-Tsoory & Abu-Akel, 2016), we hypothesized that OT stimulation would enhance social safety learning in a *trust acquisition* condition, but block (PE-enhanced) learning in a *trust violation* condition (Fig. 1). This is the first study of effects of systemic CL on social behavior, but we hypothesized that CL would not affect *trust acquisition* and facilitate (PE-enhanced) *trust violation* learning (Fig. 1). In addition, we tested the effect of OT manipulation on more general abilities (such as exploration and anxiety-like behavior, sociability and social memory, spatial working memory and contextual learning) to investigate the ability of OT to modulate any kind of (acquired) behavior. Trust-like behavior (social safety learning) was proxied by an adaptation of our social transmission of food preference (STFP) protocol (Van der Jeugd & D’Hooge, 2018) which is based on the observation that mice display neophobia when encountering novel food, but will consume it when a conspecific signals the safety of the food. We used safe food preference as a homologue of trust acquisition (STFP1), and modeled trust violation using lithium chloride (LiCl)-induced food aversion after social interaction (STFP2).

**Fig. 1.**
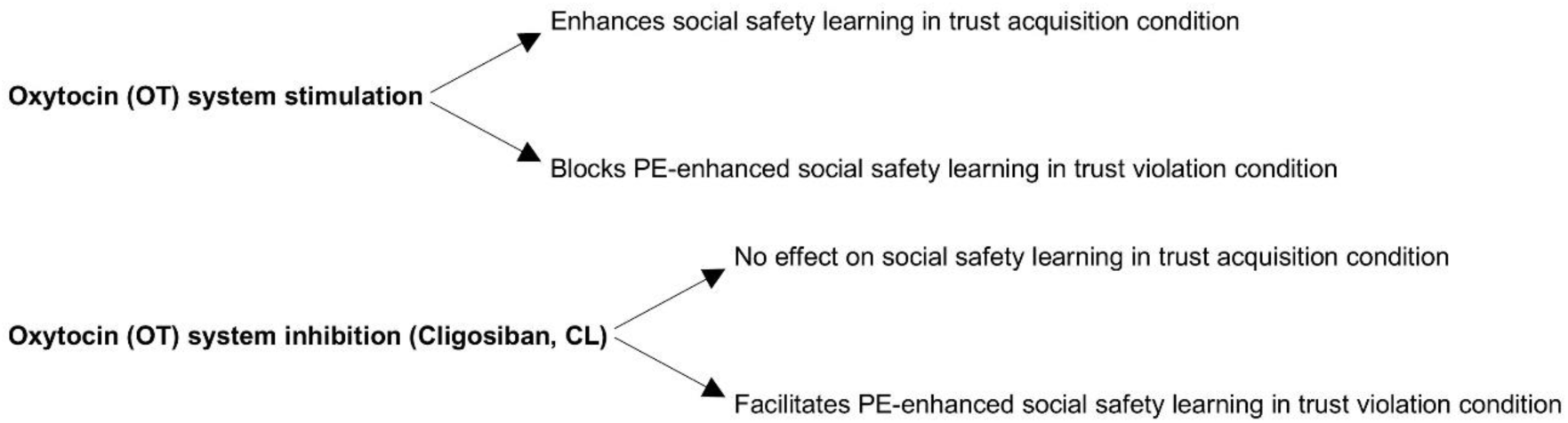
Hypothesized effects of OT system stimulation and inhibition on social safety learning in the trust acquisition and violation conditions. (PE = prediction error.)

## Materials and methods

### Animals

We used 136 C57BL/6J mice (88 male, 48 female; Janvier Labs, France), aged 10-12 weeks upon arrival, to conduct three separate experiments with different groups of mice (see Table 1 for animal numbers and sex). In **Experiments 1** and **2a,** we compared the effects of OT and CL to vehicle (VEH), respectively phosphate-buffered saline (PBS) for OT and dimethyl sulfoxide (DMSO) for CL, in males and females on explorative and anxiety-like behavior, sociability and social memory, spatial working memory, trust-like behavior and contextual associative learning. We repeated **Experiment 2a** to compare the effects of CL to VEH in males on explorative and anxiety-like behavior, sociability and social memory, and trust-like behavior in males (**Experiment 2b**). These data were merged with male data from **Experiment 2a**. In **Experiment 3**, we evaluated the effects of CL compared to VEH in males on conditioned taste aversion (CTA); (1) CTA to a liquid substance (CTA_liquid_), (2) CTA to food (CTA_food_), and sociability. During all experiments, mice lived in same-sex groups of 4 (males) to 6 (females) mice. They were housed in standard conditions (22-25 °C, humidity 50-70%, on 12h light/dark cycle with lights on at 07:00 am) in Macrolon cages containing wood-shaving bedding and cage enrichment (e.g., nesting material and toilet paper rolls to hide) with food and water available ad libitum unless otherwise specified below.

**Table 1.** Sample characteristics of the different experiments.

| Experiment | Group | Males | Females |
| --- | --- | --- | --- |
| 1. Effect of OT | OT | 12 | 12 |
|  | Vehicle control (PBS) | 12 | 12 |
| 2. Effect of CL | CL | 24 | 12 |
|  | Vehicle control (DMSO) | 24 | 12 |
| 2a. Males + females | CL | 12 | 12 |
|  | Vehicle control (DMSO) | 12 | 12 |
| 2b. Males | CL | 12 |  |
|  | Vehicle control (DMSO) | 12 |  |
| 3. CL in conditioned taste aversion | CL | 8 |  |
|  | Vehicle control (DMSO) | 8 |  |

*Note.* OT = oxytocin, PBS = phosphate-buffered saline, CL = cligosiban, DMSO = dimethyl sulfoxide.

### Preparations and administrations

All solutions were prepared as stocks, and stored in small aliquots at -20°C. Aliquots were coded to allow blinded experiments. OT, CL, and their respective VEH were administered 30 min before each behavioral test, and scheduled at least 24 h apart. OT was dissolved in PBS (1 µg/µL) and OT or PBS was administered intranasally (2 x 6 µL) using a micropipette (Neumann et al., 2013). To ensure inhalation, we carefully distributed droplets of the respective solutions over the nostrils of mice and waited for inhalation before proceeding with the rest of the solution. CL was dissolved in DMSO (10mg/mL in DMSO). This stock solution was diluted 100x with saline (SAL), VEH was 1% DMSO in SAL. CL (1mg/kg) or VEH was administered intraperitoneally (i.p.; 1% body weight). LiCl (1 mEq/kg) was dissolved in SAL and administered i.p. (1% body weight) 30 min after CTA tests (STFP2, CTA_liquid_, CTA_food_). We established this low-dose LiCl CTA protocol based on our own observations that it effectively induced taste aversion learning during CTA_liquid_ (see manipulation check in Supplementary file 1).

### Behavioral testing

The test battery varied over the different experiments but the order of tests was identical. In Experiments 1 and 2a, mouse behavior was evaluated in open field (OF), social proximity (SP), social preference/social novelty (SPSN), T-maze, STFP1 and STFP2, and passive avoidance (PA). In Experiment 2b, we included OF, SPSN, STFP1 and STFP2 in our test battery (data from these tests were merged with the male data from Experiment 2a). In Experiment 3, we evaluated mouse behavior in CTA_liquid,_ CTA_food_ and social exploration (SE). Procedures and results of control measures (OF, SP, SPSN, T-maze, PA and SE) are reported in Supplementary file 1.

Thirty minutes before every test, mice were brought to the testing room for habituation. In social tests, we assessed behavior of experimental mice towards unfamiliar mice matched in sex and strain (i.e., strangers), as well as age in STFP1 and STFP2 (i.e., demonstrators). Additionally, mice that had been tested were kept separate from naïve cage mates to avoid transmission of the scent of an unfamiliar mouse. In tasks that involved different food types (STFP1, STFP2, CTA_food_), gloves were changed to avoid contamination. Lastly, setups were always thoroughly cleaned with 70% ethanol in between mice. All procedures and behavioral testing were conducted during the light phase of mice’s cycle and the experimenter was blind to treatment throughout all testing and subsequent scoring. The procedures in the current study were approved by the Animal Ethics Committee of the University of Leuven according to the EU directive 2010/63/EU for animal experiments.

#### Social safety learning conditions

We adapted the social transmission of food preference (STFP) protocol for our conditions (Van der Jeugd & D’Hooge, 2018). The tests were conducted in a three-compartment setup that consisted of a central chamber (37.5 long, 26.5 cm wide) and 2 side chambers (27.5 cm long, 26.5 cm wide) of transparent Plexiglass, separated by sliding doors. In each side chamber, a paper cup (Ø = 4 cm) was placed in a metal holder consisting of a ramp (10 x 10 cm, height 7 cm) that allowed easy access to the food. The STFP protocol consisted of 3 phases over 5 days (*habituation*, *social interaction* and *test*). Until the end of the protocol, experimental mice (observers) had scheduled access to regular food pellets (1h/day), whereas demonstrators were put on a flavored diet by mixing flavored powder with crushed food pellets (1% w/w). Body weight of observers was monitored twice per day (before and after feeding).

In the **trust acquisition** condition (**STFP1**; Fig. 2A), observers and demonstrators were placed on a scheduled feeding regimen for two days. On day 3, observers were habituated to the setup by placing them in the central chamber of the setup for 10 min without, and 10 min with access to the side chambers in which they could find food cups with regular food pellets (*habituation* phase). On day 4, demonstrators were placed in a round wire cage (Ø = 10 cm) in the center of the central chamber to promote social interaction with the observers for 30 min (*social interaction* phase). Observers received their treatment 30 min prior to this phase. On day 5, in the *test* phase, observers could freely explore the entire setup for 2 h with access to food that was signaled as safe (i.e., paprika) in one side chamber and novel food in the other (i.e., oregano). To minimize the risk of contamination within the cages, the safe and novel food flavors were not counterbalanced.

**Fig. 2.**
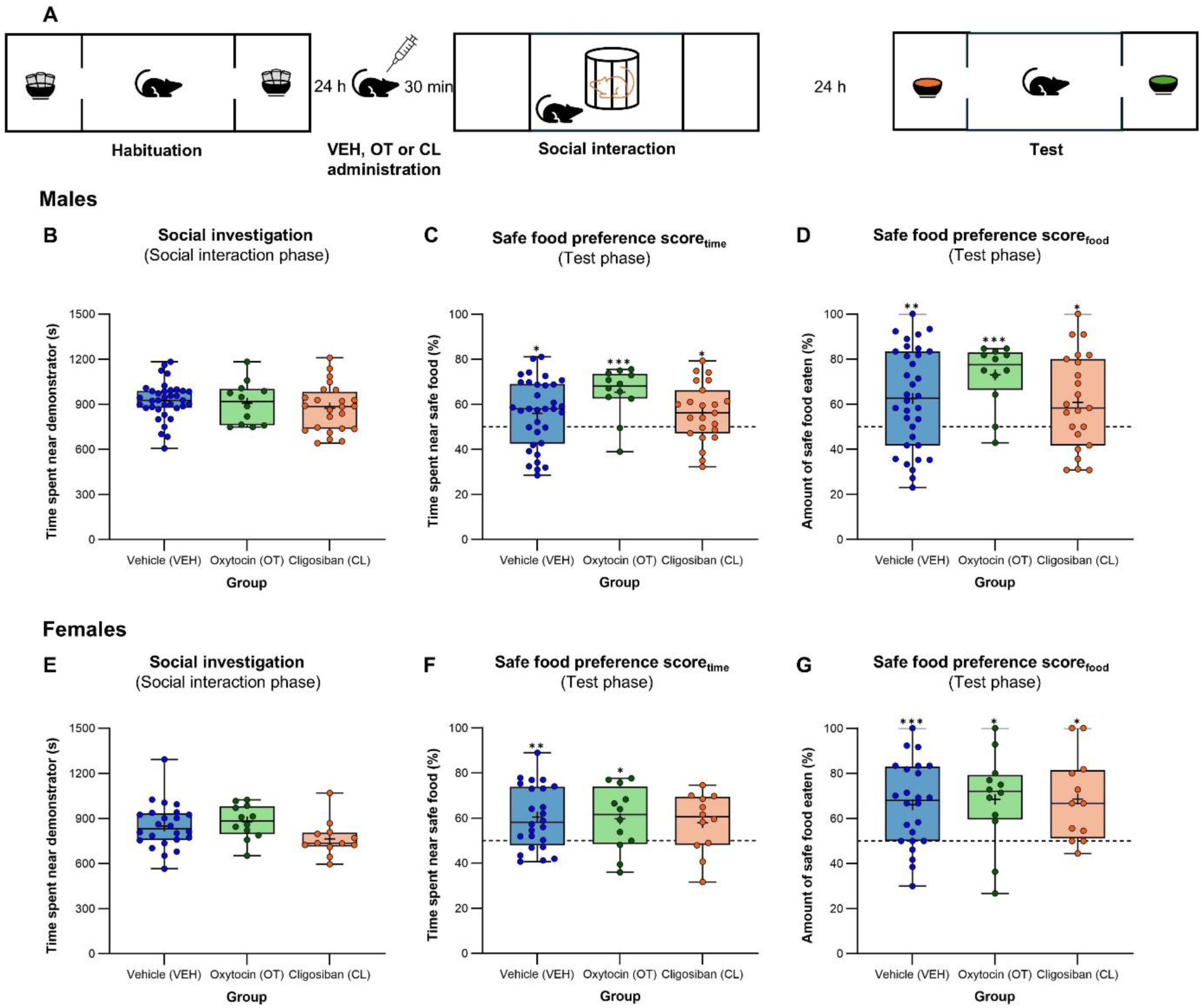
Oxytocin enhances learning in the trust acquisition condition. **(A)** Overview of the trust acquisition condition. In the *habituation* phase, the observer mouse was presented regular food (grey pellets). 30 min before the *social interaction* phase, the observer mouse received its treatment. In the *social interaction* phase, the demonstrator mouse (inside circular cage) presented a safe food (e.g., paprika). In the *test* phase, the observer mouse was presented a safe (orange = paprika-flavored) and novel food (green = oregano-flavored), respectively. **(B)** In the *social interaction* phase, VEH-treated males spent a similar amount of time near the demonstrator as those in the OT and CL group. **(C-D)** In the *test* phase, safe food preference scores of males treated with VEH, OT or CL were significantly above chance indicating that they preferred to spend time near, and eat from the safe food. Notably, safe food preference scores tended to be higher in the OT group compared to VEH-treated males. **(E)** In the *social interaction* phase, VEH-treated females spent a similar amount of time near the demonstrator as those in the OT and CL group. **(F-G)** In the *test* phase, safe food preference scores of females treated with VEH, OT or CL were (significantly) above chance without group differences. All female groups preferred to spend time near, and eat from the safe food to a similar extent. Asterisks on top of boxplots represent statistical significance of safe food preference scores compared to chance level (\**p* < .05, \*\**p* < .01, \*\*\**p* < .001).

Movements of observers were tracked by an overhead camera connected to the ANY-Maze Video Tracking System (Stoelting, Dublin, Ireland) to assess time spent in the different chambers. Additionally, before and after the *test* phase, food cups were weighed to measure the amount of food eaten. In the *social interaction* phase, we analyzed time spent near the demonstrator. In the *test* phase, trust-like behavior was measured by calculating a safe food preference score in terms of amount of safe food eaten (safe food preference score_food_ = amount of safe food eaten/(safe + novel food eaten)*100) and time spent near the safe food (safe food preference score_time_ = time spent near safe food/(time spent near safe + novel food)*100).

To model **trust violation** (**STFP2**; Fig. 3A), we repeated the STFP1 protocol but administered LiCl to observers 30 min after the *social interaction* phase. LiCl administration causes nausea and is a standard protocol to induce taste aversion, where mice learn to associate a novel taste (conditional stimulus, CS) with aversive consequences of an unconditional stimulus (UCS; e.g., LiCl), resulting in avoidant behavior towards the CS (Chambers, 2018; Welzl et al., 2001). In STFP, mice encounter a novel taste which they learn to perceive as safe through social interaction (trust-like behavior). However, pairing the taste with LiCl would replace the positive association with the novel taste with a negative association (i.e., PE; trust violation) which should result in decreased safe food preference. To avoid overlap between the **trust acquisition** and **trust violation** conditions, we used different food flavors (safe: rosemary, novel: basil) and ensured that observers were paired with different demonstrators in the *social interaction* phase. Variables of interest were similar as in the **trust acquisition** condition.

**Fig. 3.**
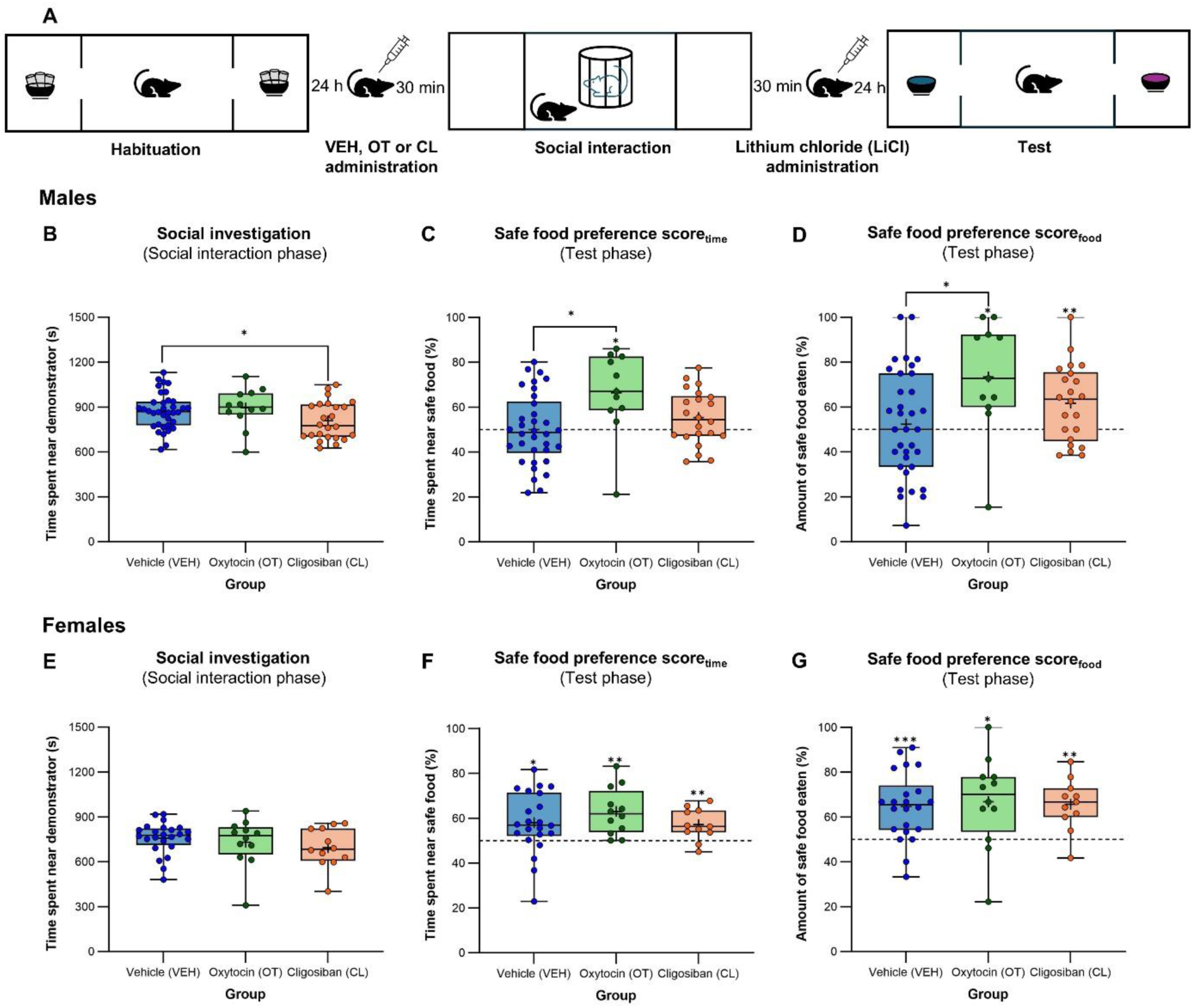
Oxytocin and cligosiban block learning in the trust violation condition. **(A)** Overview of the trust violation condition. In the *habituation* phase, the observer mouse was presented regular food (grey pellets). 30 min before the *social interaction* phase, the observer mouse received its treatment. In the *social interaction* phase, the demonstrator mouse (inside circular cage) presented a safe food (e.g., rosemary). 30 min after the *social interaction* phase, the observer mouse received a LiCl injection to induce nausea. In the *test* phase, the observer mouse was presented a safe (blue = rosemary-flavored) and novel food (purple = basil-flavored), respectively. **(B)** In the *social interaction* phase, time spent near the demonstrator before experiencing LiCl-induced nausea was similar for OT- and VEH-treated males, but slightly reduced for CL-treated males. **(C-D)** In the *test* phase, males treated with OT or CL retained safe food preference scores significantly above chance while this was not the case for VEH-treated males. Notably, safe food preference scores for OT-treated males were again enhanced compared to VEH-treated males. **(E)** In the *social interaction* phase, VEH-treated females spent a similar amount of time near the demonstrator as those in the OT or CL group. **(F-G)** In the *test* phase, females treated with VEH, OT or CL retained safe food preference scores significantly above chance without differences between groups. This indicated that all female groups retained the preference to spend time near, and eat from the safe food to a similar extent. Asterisks on top of boxplots represent statistical significance of safe food preference scores compared to chance level (\**p* < .05, \*\**p* < .01, \*\*\**p* < .001).

#### Conditioned taste aversion (CTA)

CTA tests assess the ability of mice to learn to associate a novel taste (CS) with the negative consequences of an UCS (here nausea after LiCl administration). In contrast to the trust violation condition, CTA requires mice to learn about a novel taste through direct exposure. If CTA is successful, this should result in avoidant behavior towards the novel taste. In the **CTA_liquid_** (Fig. 4A) and **CTA_food_** (Fig. 4D) protocols, we respectively used sodium saccharin (an artificial sweetener), and rosemary-flavored food as novel tastes. Taste aversion learning would be indicated by a reduced preference for the sodium saccharin solution (0.5% in tap water) over water in CTA_liquid_, and for the rosemary-flavored food over a food taste never encountered before (basil) in CTA_food_.

**Fig. 4.**
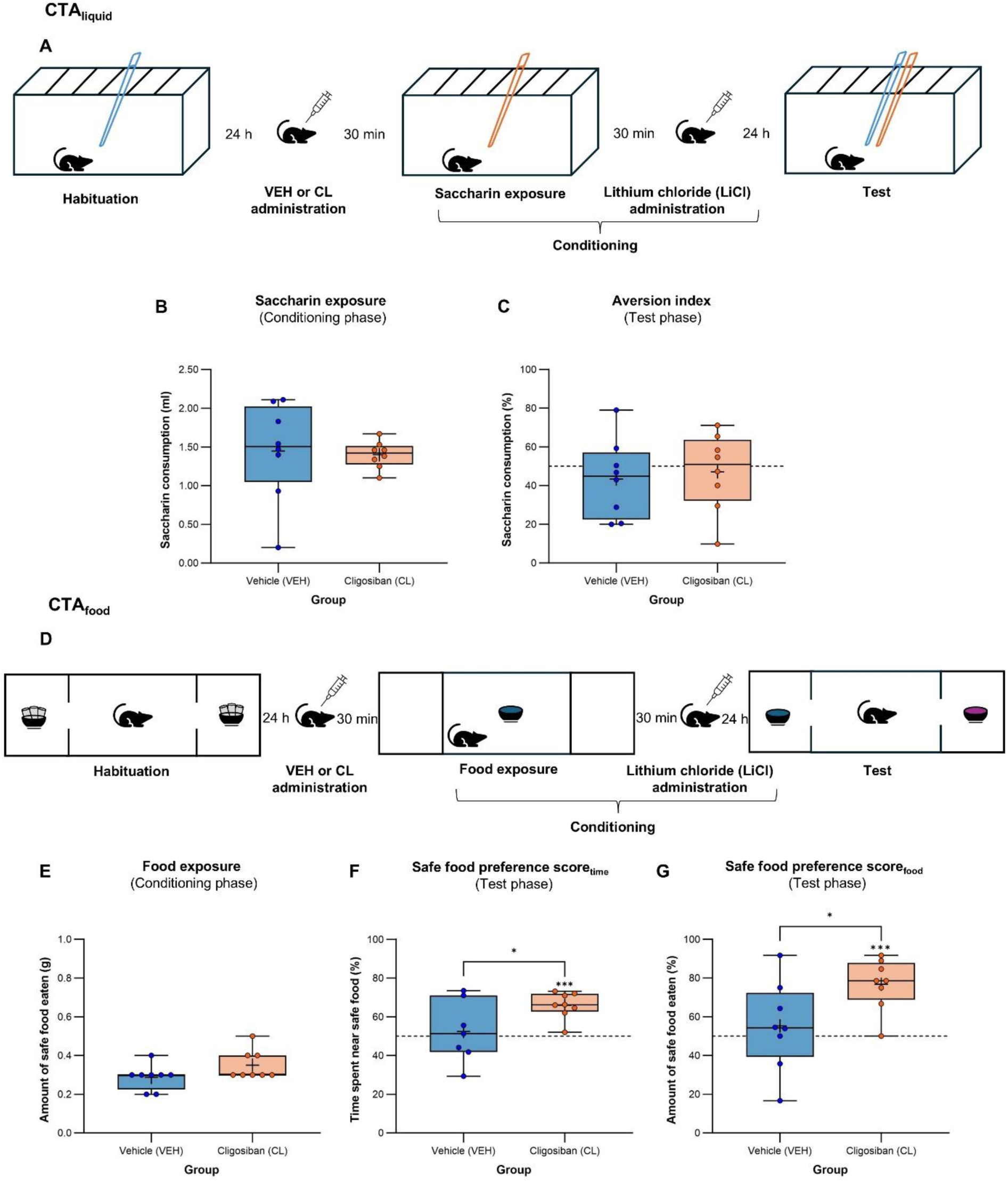
Cligosiban blocks non-social, olfactory-dependent taste aversion learning. **(A)** Overview of the CTA_liquid_ protocol. Blue and orange Falcon tubes represent the water and sodium saccharin solution, respectively. **(B)** In the *conditioning* phase of CTA_liquid_, VEH- and CL-treated mice consumed a similar amount of the sodium saccharin solution before experiencing LiCl-induced nausea. **(C)** In the *test* phase of CTA_liquid_, the AI of VEH- and CL-treated mice was not significantly above chance and did not differ between groups indicating that taste aversion learning occurred. **(D)** Overview of the CTA_food_ protocol. In the *habituation* phase, grey pellets represent regular food. In the *conditioning* and *test* phase, blue and purple crushed food pellets represent rosemary- (safe food) and basil-flavored food (novel food), respectively (same flavors that were used in the trust violation condition). **(E)** In the *conditioning* phase of CTA_food_, VEH- and CL-treated mice consumed similar amounts of food before experiencing LiCl-induced nausea. **(F-G)** In the *test* phase of CTA_food_, safe food preference scores of CL-treated mice were significantly above chance while this was not the case for VEH-treated mice. Safe food preference scores of CL-treated mice were also significantly higher compared to the VEH group. These results indicate that taste aversion learning did occur in VEH-treated mice but was blocked in the CL group. Asterisks on top of boxplots represent statistical significance of safe food preference scores compared to chance level (\**p* < .05, \*\*\**p* < .001).

The **CTA_liquid_** protocol was conducted in separate Macrolon type II test cages for each mouse. During testing, liquids were offered in 15ml polypropylene Falcon tubes, allowing for consumption measurement by weighing the tubes before and after testing. The protocol consisted of 3 phases over 5 days (*habituation*, *conditioning* and *test*). Throughout, mice had restricted access to water (30 minutes/day in their home cage). During the first 3 days, tap water was offered in Falcon tubes for 30 min each day to habituate mice to drinking from Falcon tubes (*habituation* phase). On day 4, mice were offered the sodium saccharin solution for 30 min, followed by a LiCl-injection 30 min later (*conditioning* phase). On day 5, Falcon tubes containing either water or the sodium saccharin solution were offered simultaneously and placed closely alongside each other (*test* phase) for 30 min. We analyzed sodium saccharin solution consumption in the *conditioning* phase on day 4 and calculated an aversion index (AI) regarding liquid consumption in the *test* phase on day 5 (AI = water consumption day 5/(water + sodium saccharin solution consumption day 5) x 100). A higher (more water consumption) AI indicates more taste aversion.

The **CTA_food_** protocol was conducted in accordance with the STFP2 protocol (trust violation condition) using the same setup. As indicated above, instead of learning about the safety of novel food through interaction with a demonstrator in the STFP2 protocol, mice in the CTA_food_ protocol learned about a novel taste (i.e., rosemary, safe food) through direct exposure (*conditioning* phase). Mice used in CTA_food_ were different than those used in STFP2 to ensure the novelty of the rosemary-flavored food. Other phases of the CTA_food_ protocol were not different compared to those in the STFP2 protocol, and the variables of interest were the same.

### Statistical analysis

In all experiments, data from males and females were analyzed separately by comparing group means with independent *t*-tests. Violations of the normality or homogeneity assumption were addressed by conducting the Wilcoxon rank-sum and Welch’s t-test, respectively. Safe food preference scores and the AI were compared to chance (50%) using one-sample *t*-tests. Scores significantly above or below chance were interpreted as a significantly increased or decreased preference, respectively. Since we used two outcome variables (safe food preference score_food_ and safe food preference score_time_) to assess social safety learning and CTA_food_, we corrected for multiple testing by applying the Benjamini-Hochberg method (Benjamini & Hochberg, 1995; Yekutieli & Benjamini, 1999). This correction was applied both when comparing safe food preference scores between groups and when comparing these scores to chance within each group. Cohen’s *d* was used as an estimate of the effect size for *t*-tests, and *r* for the Wilcoxon rank-sum test. Effects were considered significant at α = 0.05 and the sample sizes in the different experiments were chosen to obtain a power of 0.80.

Data is visualized below as boxplot diagrams in which all individual datapoints, median (horizontal line in box), mean (+), interquartile range (IQR) including the 25th percentile (Q1; lower end of box) and 75th percentile (Q3; upper end of box) are indicated. Visualizations include all data but if extreme outliers (Q1 – 3 x IQR or Q3 + 3 x IQR) were present, winsorizing was used to reduce the impact of single data points before conducting analyses. All analyses were conducted in R (R Core Team, 2024) and GraphPad Prism.

## Results

We merged VEH groups to increase sample size (Supplementary file 2) when differences between VEH groups of Experiment 1 (PBS) and 2 (DMSO) were not significant. Results of control measures (OF, SP, SPSN, T-maze and PA) indicated that explorative and anxiety-like behavior, sociability and social memory, spatial working memory and contextual learning were similar between groups in both males and females (Supplementary file 1).

### Oxytocin enhances learning in the trust acquisition condition (STFP1)

In the *test* phase of the trust acquisition condition, three male mice (2 VEH, 1 CL) that mixed safe and novel food (by removing their food cups from the holders) were excluded from analyses. In the *social interaction* phase, VEH-treated males (Fig. 2B) and females (Fig. 2E) spent a similar amount of time near the demonstrator as the OT (male: *t*_46_ = 0.39, *p* = .70, *d* = 0.13; female: *t*_34_ = 0.58, *p* = .57, *d* = 0.20) or CL mice (male: *t*_58_ = 1.35, *p* = .18, *d* = 0.36; female: *t*_34_ = 1.78, *p* = .08, *d* = 0.63). The following day, in the *test* phase, safe food preference scores of males (Fig. 2C-D) treated with VEH (time: *t*_33_ = 2.32, *p* = .03, *d* = 0.40; food: *t*_33_ = 3.30, *p* = .005, *d* = 0.57), OT (time: *t*_11_ = 4.82, *p* < .001, *d* = 1.39; food: *t*_11_ = 5.76, *p* < .001, *d* = 1.66) or CL (time: *t*_22_ = 2.36, *p* = .03, *d* = 0.49; food: *t*_22_ = 2.47, *p* = .03, *d* = 0.52) were significantly above chance indicating that they preferred to spend time near, and eat from the safe food. Notably, social safety learning was enhanced in the OT male group, as safe food preference scores tended to be higher compared to VEH-treated males (time: *t*_44_ = 1.97, *p* = .07, *d* = 0.66; food: *t*_31.40_ = 1.90, *p* = .07, *d* = 0.57). Safe food preference scores of females (Fig. 2F-G) treated with VEH (time: *t*_23_ = 3.64, *p* = .001, *d* = 0.74; food: *t*_23_ = 4.21, *p* < .001, *d* = 0.86), OT (time: *t*_11_ = 2.32, *p* = .04, *d* = 0.67; food: *t*_11_ = 3.07, *p* = .02, *d* = 0.89) or CL (time: *t*_11_ = 2.10, *p* = .06, *d* = 0.61; food: *t*_11_ = 3.39, *p* = .01, *d* = 0.98) were (significantly) above chance without group differences.

### Oxytocin and cligosiban block learning in the trust violation condition (STFP2)

In the *test* phase of the trust violation condition, five males (2 VEH, 1 OT, 2 CL) and three females (2 VEH, 1 CL) that mixed safe and novel food were excluded. In the *social interaction* phase, time spent near the demonstrator, before experiencing LiCl-induced nausea, was similar in all female groups (VEH-OT: *w* = 138*, p* = .86*, r* = .03; VEH-CL: *w* = 104*, p* = .19*, r* = .22; Fig. 3E) and OT-treated males (*t*_46_ = 0.53, *p* = .60, *d* = 0.18), but slightly reduced in males in the CL group (*t*_58_ = -2.03, *p* = .047, *d* = 0.53; Fig. 3B). The following day, in the *test* phase, males (Fig. 3C-D) in the OT (time: *t*_10_ = 2.91, *p* = .02, *d* = 0.88; food: *t*_10_ = 3.08, *p* = .02, *d* = 0.93) and CL group (time: *t*_21_ = 2.06, *p* = 0.05, *d* = 0.44; food: *t*_21_ = 3.18, *p* = .008, *d* = 0.68) displayed safe food preference scores significantly above chance, in contrast to VEH-treated males (time: *t*_33_ = 0.01, *p* = .99, *d* = 0.002; food: *t*_33_ = 0.58, *p* = .99, *d* = 0.10). Social safety learning was again enhanced in the OT male group, indicated by significantly higher safe food preference scores compared to VEH-treated males (time: *t*_43_ = 2.87, *p* = .01, *d* = 1; food: *t*_43_ = 2.51, *p* = .02, *d* = 0.87). Females (Fig. 3F-G) treated with VEH (time: *t*_21_ = 2.73, *p* = .01, *d* = 0.58; food: *t*_21_ = 4.52, *p* < .001, *d* = 0.96), OT (time: *t*_11_ =4.28, *p* = .003, *d* = 1.24; food: *t*_11_ = 2.87, *p* = .02, *d* = 0.83) or CL (time: *t*_10_ = 3.33, *p* = .01, *d* = 1; food: *t*_10_ = 4.45, *p* = .002, *d* = 1.34) displayed safe food preference scores significantly above chance without group differences.

### Cligosiban blocks non-social, olfactory-dependent taste aversion learning

Contrary to our hypothesis, CL-treated males displayed safe food preference in the trust violation condition, despite LiCl-induced nausea. We conducted an additional experiment (Exp. 3) with 16 male mice (8 VEH, 8 CL) to investigate whether this effect would also occur in non-social learning (CTA_food_) and non-olfactory-dependent learning (CTA_liquid_). For consistency, we used the same food flavors in CTA_food_ as in the trust violation condition (safe: rosemary, novel: basil). We included a social exploration test as a control measure, which failed to indicate differences in sociability (Supplementary file 1).

In the *conditioning* phase of the CTA_liquid_ test (Fig. 4B), VEH- and CL-treated mice consumed a similar amount of the sodium saccharin solution before experiencing LiCl-induced nausea (*t*_14_ = 0.20, *p* = .84, *d* = 0.10), indicating that CL did not alter drinking behavior. The following day, in the *test* phase (Fig. 4C), the AI of mice treated with VEH (*t*_7_ = 0.92, *p* = .39, *d* = 0.32) and CL (*t*_7_ = 0.42, *p* = .69, *d* = 0.15) did not significantly differ from chance and did not differ between groups (*t*_14_ = 0.35, *p* = .73, *d* = 0.18). This indicated that both groups associated the LiCl-induced nausea with the sodium saccharin solution, leading to a similar reduction in sodium saccharin preference, indicating similar CTA.

In the *test* phase of the CTA_food_ test, one mouse moved its food cup outside of the designated zone by the tracking system. Therefore, time spent near the safe food was not measured correctly and we excluded this mouse from this analysis (7 DMSO, 8 CL). Because there was no spillage during the test, we were still able to calculate its safe food preference score in terms of amount of food eaten. In the *conditioning* phase (Fig. 4E), VEH- and CL-treated mice consumed a similar amount of food before induction of LiCl-induced taste aversion (*t*_14_ = 1.95, *p* = .07, *d* = 0.97), indicating that CL did not affect feeding behavior. The following day, in the *test* phase (Fig. 4F-G), VEH-treated mice avoided food paired with LiCl as safe food preference scores were similar to chance (time: *t*_6_ = 0.39, *p* = .71, *d* = 0.15; food: *t*_7_ = 0.64, *p* = .71, *d* = 0.23), indicating successful induction of CTA. In contrast, CL-treated mice retained interest in food that had been paired with LiCl as safe food preference scores were significantly above chance (time: *t*_7_ = 6.58, *p* < .001, *d* = 2.33; food: *t*_7_ = 5.64, *p* < .001, *d* = 1.99). This interest was also significantly enhanced compared to the VEH group (time: *t*_13_ = 2.21, *p* = .046, *d* = 1.14; food: *t*_14_ = 2.29, *p* = .046, *d* = 1.14), indicating that CL blocked taste aversion learning.

## Discussion

We investigated social safety learning in the STFP paradigm, as a murine homologue of acquired trust. This interpretation of the paradigm is supported by research that shows that STFP acquisition is influenced by demonstrator and observer characteristics (Andrade et al., 2023; Galef et al., 1991). It reflects natural rodent behavior and requires only brief observer-demonstrator interaction (Galef & Stein, 1985). STFP acquisition depends on oronasal investigation of the demonstrator (Andrade et al., 2023). We used time spent in proximity of the demonstrator as a measure of social investigation, whereas some authors used other measures. Andrade et al. (2023) showed that only observers that had engaged in oronasal contact with the demonstrator above a defined threshold, developed a safe food preference. As a side, it would be interesting to investigate whether OT manipulation influences this threshold.

Firstly, we confirmed the hypothesis that OT enhances trust acquisition and blocks learning from trust violation (in males, but not in females). This could be attributed to the putative involvement of OT in processing of social information during the interaction between observer and demonstrator. The presence of carbon disulfide (a semiochemical on the breath of the demonstrator) is required for STFP to occur (Galef et al., 1988). The neural signals generated by food odor and CS_2_ are integrated in the olfactory bulb, produce socially-relevant olfactory input to the amygdala, and eventually form hippocampus-dependent memories about food safety (Choleris et al., 2009; Munger et al., 2010).

Hegoburu et al. (2024) identified an OT-dependent mechanism in rats whereby CeA neurons shift a stimulus’s meaning from fearful to safe. Novel food triggers a neophobic amygdala response (Lin et al., 2012), whereas during STFP, the initially neophobic response shifts and consumption of safe food is evoked. In keeping with the social salience hypothesis (Shamay-Tsoory & Abu-Akel, 2016). OT might have enhanced the salience of social information about food safety (Oettl et al., 2016), thus potentiating the safety shift mechanism and subsequent trust acquisition. Conversely, the blockage of trust violation learning by OT could also be attributed to the enhanced social salience mechanism, reducing the net impact of LiCl-induced nausea.

After receiving the selective OT antagonist CL (Wayman et al., 2018), our mice still displayed trust-like behavior. Contrary to our prediction, CL blocked learning from trust violation as well as non-social olfactory learning. This could simply be attributed to the reportedly non-linear and topical involvement of OT in (social) behavior and learning (e.g., Baldi et al., 2021; Chini et al., 2014; Popik et al., 1992). For instance, OT agonists and antagonists administration in rats enhanced or blocked fear extinction, depending on the targeted brain region (Campbell-Smith et al., 2015). Also, different effects of the selective OT agonist TGOT and OT itself on social behavior and fear extinction in rats suggest that OT interacts with other neurochemical systems (Lahoud & Maroun, 2013; Suraev et al., 2014). For instance, OT influences social behavior by binding to receptors for the structurally similar arginine-vasopressin (AVP) peptide (Huang et al., 2014; Song & Albers, 2018; Song et al., 2014; Bous et al., 2023). Notably, AVP facilitates social recognition by increasing salience of olfactory information (Rigney et al., 2022), and improves performance in difficult versions of the STFP paradigm (Choleris et al., 2009).

Manipulating the OT system did not affect performance of females in the trust acquisition and violation conditions. This can be likely attributed to differences between male and female rodent social behavior (e.g., Baumgartner et al., 2023; Misiołek et al., 2023). Females might be more attentive to social information than males (i.e., difference in social salience), reducing the impact of LiCl-induced nausea. Also, this difference might influence modulation by the OT system (Shamay-Tsoory & Abu-Akel, 2016). Moreover, male mice have higher OTR densities than females which could make them more sensitive to exogenous OT (Dumais & Veenema, 2016).

In conclusion, our results demonstrate that OT enhanced trust-like behavior and blocked learning from trust violation, possibly by influencing saliency and/or PE-driven learning. Effects of the OT antagonist not mirroring those of OT itself are possibly attributable to the involvement of other neurochemical systems, or fractal involvement of the OT system in social safety learning. We also confirmed the involvement of the OT system in olfactory-dependent tasks. Notably, none of our manipulations affected performance in females. Given that safety learning is involved in various psychiatric conditions, the OT system has been proposed as therapeutic target (Eckstein et al., 2019). Our study further emphasizes the need to consider all complexities before manipulating the system therapeutically.

## Author contributions

Samuel Budniok: Investigation, Data Curation, Visualization, Formal analysis, Writing - Original Draft, Writing - Review & Editing.

Zsuzsanna Callaerts-Vegh: Conceptualization, Formal analysis, Writing - Review & Editing, Supervision.

Marian Bakermans-Kranenburg: Conceptualization, Formal analysis, Writing - Review & Editing.

Guy Bosmans: Conceptualization, Formal analysis, Writing - Review & Editing, Funding acquisition.

Rudi D’Hooge: Conceptualization, Formal analysis, Writing - Review & Editing, Funding acquisition, Supervision.

## References

Andrade RS, Cerveira AM, Mathias M da L, Varela SAM. Interaction time with conspecifics induces food preference or aversion in the wild Algerian mouse. Behavioural Processes. 2023;211:104927.

Bakermans-Kranenburg MJ, Van IJzendoorn MH. Sniffing around oxytocin: Review and meta-analyses of trials in healthy and clinical groups with implications for pharmacotherapy. Transl Psychiatry. 2013;3:1–14.

Baldi E, Costa A, Rani B, Passani MB, Blandina P, Romano A, et al. Oxytocin and fear memory extinction: Possible implications for the therapy of fear disorders? Int J Mol Sci. 2021;22:10000.

Baumgartner NE, Biraud MC, Lucas EK. Sex differences in socioemotional behavior and changes in ventral hippocampal transcription across aging in C57Bl/6J mice. Neurobiol Aging. 2023;130:141–153.

Beery AK. Antisocial oxytocin: Complex effects on social behavior. Curr Opin Behav Sci. 2015;6:174–82.

Benjamini Y, Hochberg Y. Controlling the false discovery rate: A practical and powerful approach to multiple testing. J. R. Stat. Soc. Ser. B Methodol. 1995;57:289–300

Bosch OJ, Neumann ID. Both oxytocin and vasopressin are mediators of maternal care and aggression in rodents: From central release to sites of action. Horm Behav. 2012;61:293–303.

Bous J, Fouillen A, Orcel H, Granier S, Bron P, Mouillac B. Structures of the arginine-vasopressin and oxytocin receptor signaling complexes. Vitam Horm. 2023;123:67–107.

Brill-Maoz N, Maroun M. Extinction of fear is facilitated by social presence: Synergism with prefrontal oxytocin. Psychoneuroendocrinology. 2016;66:75–81.

Campbell-Smith EJ, Holmes NM, Lingawi NW, Panayi MC, Westbrook RF. Oxytocin signaling in basolateral and central amygdala nuclei differentially regulates the acquisition, expression, and extinction of context-conditioned fear in rats. Learn Mem. 2015;22:247–57.

Cardoso C, Ellenbogen MA, Serravalle L, Linnen AM. Stress-induced negative mood moderates the relation between oxytocin administration and trust: Evidence for the tend-and-befriend response to stress? Psychoneuroendocrinology. 2013;38:2800–4.

Chambers KC. Conditioned taste aversions. World J Otorhinolaryngol Head Neck Surg. 2018;4:92–100.

Chini B, Leonzino M, Braida D, Sala M. Learning about oxytocin: Pharmacologic and behavioral issues. Biol Psychiatry. 2014;76:360–366.

Choleris E, Clipperton-Allen AE, Phan A, Kavaliers M. Neuroendocrinology of social information processing in rats and mice. Front Neuroendocrinol. 2009;30:442–59.

Cuyvers B, van IJzendoorn M, Bakermans-Kranenburg M, Verhaeghe J, Molenberghs G, Lafit G, et al. Oxytocin and state attachment responses to secure base support after stress in middle childhood. Attach Hum Dev. 2024;26:1–21.

Diederen KMJ, Fletcher PC. Dopamine, prediction error and beyond. Neuroscientist. 2021;27:30–46.

Dulac C, O’Connell LA, Wu Z. Neural control of maternal and paternal behaviors. Science. 2014;345:765–70.

Dumais KM, Veenema AH. Vasopressin and oxytocin receptor systems in the brain: Sex differences and sex-specific regulation of social behavior. Front Neuroendocrinol. 2016;40:1–23.

Eckstein M, Almeida de Minas AC, Scheele D, Kreuder AK, Hurlemann R, Grinevich V, et al. Oxytocin for learning calm and safety. Int J Psychophysiol. 2019;136:5–14.

Galef BG, Beck M, Whiskin EE. Protein deficiency magnifies social influence on the food choices of Norway rats (Rattus norvegicus). J. Comp. Psychol. 1991;105:55–59.

Galef BG, Mason JR, Preti G, Bean NJ. Carbon disulfide: A semiochemical mediating socially-induced diet choice in rats. Physiol Behav. 1988;42:119–24.

Galef BG, Stein M. Demonstrator influence on observer diet preference: Analyses of critical social interactions and olfactory signals. Anim Learn Behav. 1985;13:31–38.

Hegoburu C, Tang Y, Niu R, Ghosh S, Triana Del Rio R, de Araujo Salgado I, et al. Social buffering in rats reduces fear by oxytocin triggering sustained changes in central amygdala neuronal activity. Nat Commun. 2024;15:2081.

Heinrichs M, Domes G. Neuropeptides and social behaviour: Effects of oxytocin and vasopressin in humans. Prog Brain Res. 2008;170:337–50.

Huang H, Michetti C, Busnelli M, Managò F, Sannino S, Scheggia D, et al. Chronic and acute intranasal oxytocin produce divergent social effects in mice. Neuropsychopharmacology. 2014;39:1102–14.

Ide JS, Nedic S, Wong KF, Strey SL, Lawson EA, Dickerson BC, et al. Oxytocin attenuates trust as a subset of more general reinforcement learning, with altered reward circuit functional connectivity in males. Neuroimage. 2018;174:35–43.

Lahoud N, Maroun M. Oxytocinergic manipulations in corticolimbic circuit differentially affect fear acquisition and extinction. Psychoneuroendocrinology. 2013;38:2184–95.

Laing PAF, Vervliet B, Dunsmoor JE, Harrison BJ. Pavlovian safety learning: An integrative theoretical review. Psychon Bull Rev. 21 August 2024. 10.3758/s13423-024-02559-4

Lin JY, Roman C, Arthurs J, Reilly S. Taste neophobia and c-Fos expression in the rat brain. Brain Res. 2012;1448:82–88.

Loth MK, Donaldson ZR. Oxytocin, dopamine, and opioid interactions underlying pair bonding: Highlighting a potential role for microglia. Endocrinology. 2021;162:bqaa223.

Misiołek K, Klimczak M, Chrószcz M, Szumiec Ł, Bryksa A, Przyborowicz K, et al. Prosocial behavior, social reward and affective state discrimination in adult male and female mice. Sci Rep. 2023;13:5583.

Munger SD, Leinders-Zufall T, McDougall LM, Cockerham RE, Schmid A, Wandernoth P, et al. An olfactory subsystem that detects carbon disulfide and mediates food-related social learning. Curr. Biol. 2010;20:1438–44.

Neumann ID, Maloumby R, Beiderbeck DI, Lukas M, Landgraf R. Increased brain and plasma oxytocin after nasal and peripheral administration in rats and mice. Psychoneuroendocrinology. 2013;38:1985–93.

Oettl LL, Ravi N, Schneider M, Scheller MF, Schneider P, Mitre M, et al. Oxytocin enhances social recognition by modulating cortical control of early olfactory processing. Neuron. 2016;90:609–621.

Popik P, Vetulani J, van Ree JM. Low doses of oxytocin facilitate social recognition in rats. Psychopharmacology. 1992;106:71–74.

R Core Team (2024). R: A language and environment for statistical computing. R foundation for statistical Computing, Vienna, Austria. URL: https://www.R-project.org.

Rigney N, De Vries GJ, Petrulis A, Young LJ. Oxytocin, vasopressin, and social Behavior: From neural circuits to clinical opportunities. Endocrinology. 2022;163:bqac111.

Shamay-Tsoory SG, Abu-Akel A. The social salience hypothesis of oxytocin. Biol Psychiatry. 2016;79:194–202.

Shorey S, Asurlekar AR, Chua JS, Lim LHK. Influence of oxytocin on parenting behaviors and parent–child bonding: A systematic review. Dev Psychobiol. 2023;65:e22359.

Shyken JM, Petrie RH. Oxytocin to induce labor. Clin Obstet Gynecol. 1995;38:232–245.

Smith AS, Wang Z. Hypothalamic oxytocin mediates social buffering of the stress response. Biol Psychiatry. 2014;76:281–88.

Song Z, Albers HE. Cross-talk among oxytocin and arginine-vasopressin receptors: Relevance for basic and clinical studies of the brain and periphery. Front Neuroendocrinol. 2018;51:14–24.

Song Z, McCann KE, McNeill JK, Larkin TE, Huhman KL, Albers HE. Oxytocin induces social communication by activating arginine-vasopressin V1a receptors and not oxytocin receptors. Psychoneuroendocrinology. 2014;50:14–19.

Suraev AS, Bowen MT, Ali SO, Hicks C, Ramos L, McGregor IS. Adolescent exposure to oxytocin, but not the selective oxytocin receptor agonist TGOT, increases social behavior and plasma oxytocin in adulthood. Horm Behav. 2014;65:488–496.

Uvnäs-Moberg K, Ekstrom-Bergstrom A, Buckley S, Massarotti C, Pajalic Z, Luegmair K, et al. Maternal plasma levels of oxytocin during breastfeeding-a systematic review. PLoS One. 2020;15:e0235806.

Van der Jeugd A, D’Hooge R. Assessment of social transmission of food preferences behaviors. J Vis Exp. 2018;131:57029.

Wayman C, Russell R, Tang K, Weibly L, Gaboardi S, Fisher L, et al. Cligosiban, a novel brain-penetrant, selective oxytocin receptor antagonist, inhibits ejaculatory physiology in rodents. J Sex Med. 2018;15:1698–1706.

Welzl H, D’adamo P, Lipp H-P. Conditioned taste aversion as a learning and memory paradigm. Behav Brain Res. 2001;125:205–13.

Yekutieli D, Benjamini Y. Resampling-based false discovery rate controlling multiple test procedures for correlated test statistics. J. Stat. Plan. Inference. 1999;82:171–96.

