## Supplementary file 1 for "Oxytocin enhances acquisition in a social trust task in mice, whereas both oxytocin and its antagonist block trust violation learning"

**Methods**

**Behavioral testing**

***Tests of explorative and anxiety-like behavior, spatial working memory and contextual learning***

**Open field (OF).** Open field exploration and anxiety-like behavior were assessed by placing dark-habituated mice in a brightly-illuminated Plexiglass arena (50 x 50 cm) enclosed in a cupboard for 10 minutes. Throughout the test, mouse movements were recorded by an overhead camera connected to the ANY-Maze Video Tracking System (Stoelting, Dublin, Ireland). We measured explorative behavior by analyzing path length (in m). Anxiety-like behavior was indicated by time spent in the center and periphery (thigmotaxis), respectively. Time spent in the center is considered reduced anxiety-like behavior in rodents (Bollen et al., 2015).

**Spatial working memory T-maze.** The T-maze test taps into spatial working memory abilities. The maze consisted of 3 arms made of dark grey plastic arranged in a T, with 2 goal arms (47 cm long, 6 cm wide, enclosed by 4 cm walls) and one start arm (50 cm long, 6 cm wide, enclosed by 4 cm walls), placed on legs to elevate the maze (40 cm above the table). The goal arms could be blocked by a barrier (10 cm within arm). Mouse movements were recorded by an overhead camera connected to a tracking software (Ethovision, Noldus, Wageningen).

The protocol consisted of 2 consecutive trials of 10 min. During the first trial, a mouse was placed in the start arm and only one goal arm was accessible (familiar arm; the position of the familiar arm was counterbalanced between groups). Mice like to explore novel environments to scavenge for food. To this end, they need to remember in which areas they have been before. After 10 min, the mouse was briefly placed back in a holding cage, the maze was cleaned thoroughly with 70% ethanol and the barrier was removed. Then the mouse was placed back in the start arm for the second trial. The novelty seeking drive will have mice preferentially explore the newly accessible arm (novel arm). In both trials, we measured path length (in m) and in the second trial, we calculated a novel arm preference score (time spent in the novel arm/(time spent in the novel + familiar arm)*100) to measure mice's ability to flexibly adapt their behavior based on information in the working memory.

**Passive avoidance (PA).** Contextual learning was assessed over 2 days using a step-through setup consisting of a small, brightly illuminated compartment and a larger dark compartment, separated by a sliding door. In the dark compartment, a grid floor was connected to a current shocker. Mice received treatment 30 minutes before testing on day 1 after which they were dark-habituated. Hereafter, they were placed in the illuminated compartment of the setup. After 5 s, a sliding door opened, allowing access to the dark compartment. When mice entered the dark compartment with all 4 paws, the sliding door closed and a foot shock was delivered (0.5 mA, 2 s) after 30 s. On day 2, we repeated this procedure but without administration of the foot shock. Contextual learning was assessed by measuring the step-through latency (in s) on day 2. If mice did not enter the dark compartment within 300 s, they were taken out of the setup.

***Tests of sociability and social memory***

**Social proximity (SP).** The SP protocol was adapted from Defensor et al. (2011). It allows for both quantitative and qualitative assessment of mice's social behavior. During this test, mice were placed in a Plexiglass cylinder (Ø = 20 cm) for 10 min to promote social interaction with a stranger mouse. A video-camera placed above the cylinder recorded the interaction. After testing, the following behaviors were scored manually by the experimenter if initiated by the experimental mouse (Defensor et al., 2011): nose-nose contact (NN), nose-head contact (NH), nose-anogenital contact (NA), upright posture (U, mice display a reared posture with nose or vibrissae contact), crawling over (CO), crawling under (CU), jump escape (JE), self-grooming (SG) and allogrooming (AG). Frontal contact behavior types (NN, NH, and U) were aggregated to obtain a frontal contact score per mouse. We analyzed the frequency of frontal contacts as well as of the remaining behavior types and the total duration (in s) engaged in SG and AG.

**Social preference/ Social novelty (SPSN).** Sociability and social memory were assessed in a three-compartment setup. The setup consisted of a central chamber (40 cm long, 10 cm wide), and 2 side chambers (10 x 10 cm) made from transparent Plexiglass that were separated by a perforated wall (1 cm diameter holes), allowing limited tactile, olfactory, visual and auditory contact. Movements of the experimental mouse were recorded by an overhead camera connected to the ANY-Maze Video Tracking System (Stoelting, Dublin, Ireland).

The mice were tested in 3 consecutive episodes. Experimental mice were placed in the central chamber. After 5 min habituation period (Episode 1), a stranger mouse (S1) was placed in one of the side chambers and approach to S1 was recorded for 10 min (Episode 2). After 10 min, a second stranger (S2) was added to the other side chamber (Episode 3) and approach to either S1 or S2 was recorded. The position of S1 was counterbalanced between groups. After Episode 3, the experimental mouse and the stranger mice were placed back in their respective home cages, and the setup was thoroughly cleaned with 70% ethanol. We analyzed path length (in m) per episode as measure of activity level and calculated a stranger preference score in terms of time spent in the different zones for both Episode 2 (time spent near S1/(time spent near S1 + near empty zone)*100) and Episode 3 (time spent near S2/(time spent near S2 + near S1)*100) to obtain a measure of respectively sociability and social memory.

**Social exploration (SE).** The SE test was conducted in the OF setup but with the addition of a round wire cage (Ø = 10 cm) containing a stranger mouse in the center to obtain a measure of sociability. Mice could explore the setup for 30 minutes while their movements were recorded by an overhead camera connected to the ANY-Maze Video Tracking System (Stoelting, Dublin, Ireland). We assessed activity level and sociability by respectively analyzing path length (in m) and calculating a stranger preference score (time spent near the stranger/(time spent near the stranger + periphery)*100). We also analyzed the stranger preference score per 10-min time bin.

**Statistical analysis**

In all experiments, data from females and males were analyzed separately. Step-through latencies on day 2 of the PA protocol were analyzed using the Kaplan-Meier survival analysis to estimate and generate survival curves for each group, which represent the proportion of mice not entering the dark box (i.e., surviving) over time. The log-rank test was then conducted to compare survival curves between groups. All other data were analyzed by comparing group means with independent *t*-tests. If the normality or homogeneity assumption was violated, we computed respectively the Wilcoxon rank-sum and Welch's t-test. Additionally, all preference scores and the AI were analyzed using one-sample *t*-tests to compare these scores to chance (50%). Scores significantly above or below chance were interpreted as a significantly increased or decreased preference, respectively.

Activity levels in SPSN and T-maze and the SE stranger preference score per 10-min time bin were analyzed with two-way mixed ANOVA with treatment group as between-subjects variable and episode (SPSN), trial (T-maze) or time bin (SE) as within-subjects variables. All interactions between variables were also analyzed. Post-hoc analyses used the Benjamini-Hochberg method to correct for multiple testing [24, 25]. This method was also used to correct for multiple testing across measures assessing sociability (behaviors in SP and the S1 preference score in SPSN), and anxiety-like behavior (time spent in center and periphery in OF) in Experiments 1 and 2. Generalized eta squared ($\hat{\eta}^2_G$) was used as an estimate of the effect size for ANOVA effects, Cohen's *d* for post-hoc comparisons and independent or one-sample *t*-tests, and *r* for Wilcoxon rank-sum test. Effects were considered significant at $\alpha = 0.05$.

### Results

#### Manipulation check

To establish a CTA protocol with a low lithium chloride (LiCl) dose (1 mEq), we conducted a manipulation check in which we administered the $CTA_{liquid}$ test to 16 male mice (8 VEH, 8 LiCl). During this test, one mouse failed to consume the sodium saccharin solution during the *conditioning* phase of the protocol hence precluding its ability to associate the solution with LiCl-induced nausea. Therefore, we excluded this mouse from analyses resulting in a sample of 15 mice (8 VEH, 7 LiCl).

In the *conditioning* phase, VEH- and LiCl-treated mice consumed a similar amount of the sodium saccharin solution ($t_{13} = 0.17$, $p = .87$, $d = 0.09$). The following day, in the *test* phase, the AI of the VEH group was significantly below chance ($t_7$ = -5.96, $p < .001$, $d$ = 2.11) indicating a preference to drink from the sodium saccharin solution over water. This preference was attenuated in the LiCl group ($t_6$ = 0.12, $p$ = .91, $d$ = 0.05). Moreover, the AI of VEH-treated mice was significantly lower compared to the AI of the LiCl group ($t_{13}$ = -2.69, $p$ = .02, $d$ = 1.39) indicating an enhanced preference to drink from the sodium saccharin solution for mice in the VEH group. Together, these results indicated that a 1 mEq LiCl dose was sufficient for taste aversion learning to occur hence we used this dose in CTA tests (STFP2, $CTA_{food}$, $CTA_{liquid}$) of all subsequent experiments.

**Effects of OT and CL (Experiment 1 and 2)**

We merged VEH groups to increase sample size when differences between VEH groups of Experiment 1 (PBS) and 2 (DMSO) were not significant. Apart from male pathlength in OF, T-maze and SPSN, results indicated no significant differences between VEH groups (Supplementary file 2). Hence, for all other analyses we used one (same-sex) VEH group. Below, we use PBS and DMSO to refer to the VEH group of respectively OT and CL and VEH to refer to the merged control group.

***Tests of explorative and anxiety-like behavior, spatial working memory and contextual learning***

**Open Field.** During the OF test, males in the PBS and DMSO group had similar pathlengths as those in respectively the OT ($w$ = 53, $p$ = .29, $r$ = .22) and CL group ($t_{46}$ = 0.53, $p$ = .60, $d$ = 0.15). Pathlength was significantly increased for OT- ($t_{34}$ = 2.19, $p$ = .04, $d$ = 0.73), but not for CL- treated females, compared to VEH-treated females ($t_{34}$ = 0.78, $p$ = .44, $d$ = 0.27). Furthermore, all male groups spent an equal amount of time in the periphery (VEH-OT: $t_{46}$ = 0.66, $p$ = .51, $d$ = 0.22; VEH-CL: $w$ = 370, $p$ = .71, $r$ = .12) and center (VEH-OT: $w$ = 247, $p$ = .51, $r$ = .11; VEH-CL: $w$ = 453, $p$ = .76, $r$ = .04). Similarly, we found no differences between VEH- and OT- or CL-treated females on time spent in the periphery (VEH-OT: $t_{34}$ = 1.21, $p$ = .37, $d$ = 0.43; VEH-CL: $t_{34}$ = 0.03, $p$ = .98, $d$ = 0.01) and the center (VEH-OT: $t_{34}$ = 0.92, $p$ = .37, $d$ = 0.35; VEH-CL: $w$ = 147, $p$ = .98, $r$ = .02). Despite increased activity levels in OT-treated females, these results indicated that explorative and anxiety-like behavior were largely similar across groups in both sexes.

**T-maze.** During the T-maze test, two mice managed to climb over the door that was used to block of the novel arm during the first trial. Because this decreased the novelty of the novel arm during the second trial, these mice were excluded from analyses (Female: 1 PBS, male: 1 OT). A two-way mixed ANOVA on male pathlength with group (PBS vs. OT or DMSO vs. CL) as between-subjects variable and trial (1 vs. 2) as within-subjects variable revealed neither an interaction between group and trial (OT: $F_{1,21}$ = 0.46, $p$ = .51, $\hat{\eta}^2_G$ = .006; CL: $F_{1,22}$ = 0.45, $p$ = .51, $\hat{\eta}^2_G$ = .004) nor main effects of group (OT: $F_{1,21}$ = 0.29, $p$ = .59, $\hat{\eta}^2_G$ = .01; CL: $F_{1,22}$ = 0.00, $p$ = .98, $\hat{\eta}^2_G$ = .00) or trial (OT: $F_{1,21}$ = 1.49, $p$ = .24, $\hat{\eta}^2_G$ = .02; CL: $F_{1,22}$ = 1.82, $p$ = .19, $\hat{\eta}^2_G$ = .02) indicating similar activity levels over groups and trials. Results were largely similar in females with neither an interaction between group and trial (OT: $F_{1,33}$ = 3.37, $p$ = .08, $\hat{\eta}^2_G$ = .01; CL: $F_{1,33}$ = 0.003, $p$ = .96, $\hat{\eta}^2_G$ = .00) nor a main effect of group (OT: $F_{1,33}$ = 0.004, $p$ = .95, $\hat{\eta}^2_G$ = .00; CL: $F_{1,33}$ = 1.45, $p$ = .24, $\hat{\eta}^2_G$ = .04). However, we found an effect of trial ($F_{1,33}$ = 13.89, $p$ < .001, $\hat{\eta}^2_G$ = .04) with OT- and VEH-treated females displaying a significantly higher pathlength during trial 1 compared to trial 2. We did not find this difference in the analysis with DMSO- and CL-treated females ($F_{1,33}$ = 2.53, $p$ = .12, $\hat{\eta}^2_G$ = .01).

Furthermore, VEH (male: $t_{23}$ = 7.37, $p$ < .001, $d$ = 1.50; female: $t_{22}$ = 3.94, $p$ < .001, $d$ = 0.82), OT (male: $t_{10}$ = 5.78, $p$ < .001, $d$ = 1.74; female: $t_{11}$ = 2.88, $p$ = .01, $d$ = 0.83) and CL (male: $t_{11}$ = 5.94, $p$ < .001, $d$ = 1.71; female: $t_{11}$ = 2.74, $p$ = .02, $d$ = 0.79) groups across both sexes had novel arm preference scores significantly above chance, indicating that all groups preferred to spend time in the novel arm during trial 2. However, there were no significant group differences between males treated with VEH and OT ($w$ = 126, $p$ = .85, $r$ = .04) or CL ($t_{34}$ = 1.19, $p$ = .24, $d$ = 0.41), and between females treated with VEH and OT ($t_{33}$ = 0.27, $p$ = .79, $d$ = 0.09) or CL ($t_{33}$ = 0.18, $p$ = .86, $d$ = 0.07) indicating similar spatial working memory abilities across groups in both sexes.

**Passive Avoidance.** Survival curves for VEH-treated males did not differ significantly from those of males in the OT ($\chi^2$ (1) = 0.02, $p$ = .88) or CL group ($\chi^2$ (1) = 0.02, $p$ = .89). Similarly, survival curves for VEH-treated females were not significantly different from those in the OT ($\chi^2$ (1) = 1.98, $p$ = .16) or CL group ($\chi^2$ (1) = 2.50, $p$ = .11). These results indicated that the proportion of mice entering the dark box (i.e., step-through latency) over time on day 2 was comparable across groups in both sexes, suggesting similar contextual learning.

***Tests of sociability and social memory***

**Social Proximity.** During the SP test, 2 female (1 DMSO, 1 CL) and 6 male (3 PBS, 1 DMSO, 1 OT, 1 CL) mice were excluded from analyses because they displayed substantial escape behavior throughout the test without any intention to interact with the stranger. Furthermore, JE (male: 14/42, female: 12/46), CO (male: 13/42, female: 8/46), CU (male: 4/42, female: 5/46) and AG (male: 19/42, female: 22/46) were rarely observed in our sample and therefore, we excluded these behavior types from analyses.

VEH-treated males engaged to a similar extent as OT- and CL-treated males in frontal (VEH-OT: $t_{29}$ = 0.61, $p$ = .71, $d$ = 0.23; VEH-CL: $t_{29}$ = 0.29, $p$ = .96, $d$ = 0.11) and NA contacts (VEH-OT: $t_{29}$ = 0.38, $p$ = .71, $d$ = 0.14; VEH-CL: $t_{29}$ = 0.05, $p$ = .96, $d$ = 0.02), and spent a similar amount of time SG (VEH-OT: $w$ = 125, $p$ = .71, $r$ = .11; VEH-CL: $w$ = 114, $p$ = .96, $r$ = .03). Similarly, VEH-treated females engaged to a similar extent as OT- and CL-treated females in frontal (VEH-OT: $t_{33}$ = 0.03, $p$ = .97, $d$ = 0.01; VEH-CL: $t_{32}$ = 1.51, $p$ = .36, $d$ = 0.56) and NA contacts (VEH-OT: $w$ = 131, $p$ = .97, $r$ = .04; VEH-CL: $w$ = 90, $p$ = .36, $r$ = 0.23), and spent a similar amount of time SG (VEH-OT: $w$ = 178.5, $p$ = .65, $r$ = .24; VEH-CL: $w$ = 143, $p$ = .74, $r$ = .10). These results indicated that sociability in SP was similar across all male and female groups.

**Social Preference/ Social Novelty.** During SPSN, male and female pathlength were analyzed using a two-way mixed ANOVA with group (male: PBS vs. OT or DMSO vs. CL; female: VEH vs. OT or VEH vs. CL) as between-subjects variable and episode (1 vs. 2 vs. 3) as within-subjects variable. Group interacted significantly with episode in the analysis with males treated with CL ($F_{1.48,68.04}$ = 4.30, $p$ = .03, $\hat{\eta}^2_G$ = .03) but not with OT ($F_{1.53,33.67}$ = 3.01, $p$ = .08, $\hat{\eta}^2_G$ = .05). Conversely, group interacted significantly with episode in the analysis with females treated with OT ($F_{2,68}$= 13.22, $p$ < .001, $\hat{\eta}^2_G$ = .08) but not with CL ($F_{2,68}$ = 1.01, $p$ = .37, $\hat{\eta}^2_G$ = .01). Post-hoc analyses on the pathlength of males treated with CL compared to DMSO did not survive controlling for multiple testing while females treated with OT had a significantly higher pathlength than those treated with VEH during Episode 1 ($t_{102}$ = 5.80, $p$ < .001, $d$ = 1.86) and Episode 3 ($t_{102}$ = 3.07, $p$ = .004, $d$ = 1.32). In absence of significant interactions, we did also not find group differences over all episodes in males treated with OT compared to PBS-treated males ($F_{1,22}$ = 2.55, $p$ = .13, $\hat{\eta}^2_G$ = .07) and females treated with CL compared to VEH-treated females ($F_{1,34}$ = 0.67, $p$ = .42, $\hat{\eta}^2_G$ = .02). Overall, these findings indicated that activity levels did not differ between male groups during all episodes. Conversely, activity levels of OT-treated females were significantly increased compared to VEH-treated females during Episode 1 and 3.

Furthermore, we found an effect of episode for both males (OT: $F_{1.53,33.67}$ = 132.60, $p$ = < .001, $\hat{\eta}^2_G$ = .70; CL: $F_{1.48,68.04}$ = 188.52, $p$ < .001, $\hat{\eta}^2_G$ = .61) and females (OT: $F_{2.68}$ = 172.25, $p$ = < .001, $\hat{\eta}^2_G$ = .54; CL: $F_{2,68}$ = 120.48, $p$ < .001, $\hat{\eta}^2_G$ = .45). Specifically, the pathlength during Episode 1 was significantly higher compared to Episode 2 in males (OT: $t_{69}$ = 9.82, $p$ < .001, $d$ = 2.91; CL: $t_{141}$ = 12.70, $p$ < .001, $d$ = 2.52) and females (OT: $t_{105}$ = 8.11, $p$ < .001, $d$ = 1.77; CL: $t_{105}$ = 8.33, $p$ < .001, $d$ = 1.83), and Episode 3 in males (OT: $t_{69}$ = 10.7, $p$ < .001, $d$ = 2.85; CL: $t_{141}$ = 12.30, $p$ < .001, $d$ = 2.34) and females (OT: $t_{105}$ = 7.40, $p$ < .001, $d$ = 1.67; CL: $t_{105}$ = 8.10, $p$ < .001, $d$ = 1.98). In line with expectations, activity decreased across groups in both sexes upon the introduction of social stimuli during the test.

In Episode 2, males and females treated with VEH (male: $t_{35}$ = 3.55, $p$ = .001, $d$ = 0.59; female: $t_{23}$ = 6.23, $p$ < .001, $d$ = 1.27), OT (male: $t_{11}$ = 2.46, $p$ = .03, $d$ = 0.71; female: $t_{11}$ = 3.22, $p$ = .01, $d$ = 0.93) and CL (male: $t_{23}$ = 5.88, $p$ < .001, $d$ = 1.20; $t_{11}$ = 2.61, $p$ = .02, $d$ = 0.75) had S1 preference scores significantly above chance indicating that they preferred to spend time near S1. We found no significant differences in S1 preference scores between males and females treated with VEH compared to those treated with OT (male: $t_{46}$ = 0.67, $p$ = .71, $d$ = 0.22; female: $t_{34}$ = 0.14, $p$ = .97, $d$ = 0.05) or CL (male: $t_{58}$ = 1.58, $p$ = .48, $d$ = 0.42; female: $t_{14.6}$ = 0.12, $p$ = .91, $d$ = 0.04). This indicated a similar preference for S1 across male and female groups, suggesting comparable sociability in SPSN.

In Episode 3, males and females treated with VEH (male: $t_{35}$ = 1.51, $p$ = .14, $d$ = 0.25; female: $t_{23}$ = 1.58, $p$ = .13, $d$ = 0.32), OT (male: $t_{11}$ = 1.08, $p$ = .30, $d$ = 0.31; female: $t_{11}$ = 0.11, $p$ = .92, $d$ = 0.03) and CL (male: $t_{23}$ = 1.15, $p$ = .26, $d$ = 0.24; female: $t_{11}$ = -2.13, $p$ = .06, $d$ = 0.62) had S2 preference scores similar to chance indicating no preference to spend time near S2. We found no significant differences in S2 preference scores between males treated with VEH and those treated with OT ($t_{46}$ = 1.84, $p$ = .07, $d$ = 0.61) or CL ($t_{58}$ = 1.87, $p$ = .07, $d$ = 0.49). S2 preference scores were also not different between females in the VEH and OT group ($t_{34}$ = 0.70, $p$ = .49, $d$ = 0.25), but significantly lower for CL-treated females compared to the VEH group ($t_{34}$ = -2.29, $p$ = .03, $d$ = 0.81). These results indicated that interest for S2 was similar across male groups but reduced for CL females, suggesting largely similar social memory.

**Cligosiban in conditioned taste aversion (Experiment 3)**

***Test of sociability***

**Social Exploration.** VEH- and CL-treated mice did not differ in pathlength during the SE test ($t_{14}$ = 0.78, $p$ = .45, $d$ = 0.39) and had similar stranger preference scores ($t_{14}$ = 0.84, $p$ = .42, $d$ = 0.42) indicating similar activity levels and similar social interest. Surprisingly, the stranger preference score of both VEH- ($t_7$ = 0.14, $p$ = .89, $d$ = 0.05) and CL-treated mice ($t_7$ = 0.93, $p$ = .39, $d$ = 0.33) was also not significantly above chance indicating that they had no preference to spend time near the stranger during the test. Given that the test lasted 30 minutes, this null result could have been caused by decreased interest in the stranger towards the end of the test.

Therefore, we analyzed the stranger preference score in 10-min time bins by conducting a two-way mixed ANOVA with group (VEH vs. CL) as between-subjects variable and time bin (1 vs. 2 vs. 3) as within-subjects variable. Group did not interact significantly with time bin ($F_{2,28}$ = 0.15, $p$ = .86, $\hat{\eta}^2_G$ = .003) and there was also no significant effect of group ($F_{1,14}$ = 0.81, $p$ = .38, $\hat{\eta}^2_G$ = .04). However, we did find a significant effect of time bin ($F_{2,28}$ = 4.71, $p$ = .02, $\hat{\eta}^2_G$ = .09) with the highest stranger preference score in the first time bin ($M$ = 55.44) which decreased over time (time bin 2: $M$ = 51.22, time bin 3: $M$ = 47.63). The stranger preference score during time bin 1 across both groups was significantly above chance ($t_{15}$ = 2.67, $p$ = .02, $d$ = 0.67) indicating that during this period mice preferred to spend time near the stranger.

**References**


Bollen B, Ramanantsoa N, Naert A, Matrot B, Van den Bergh O, D'Hooge R, et al. Emotional disorders in adult mice heterozygous for the transcription factor Phox2b. Physiol Behav. 2015;141:120–26.

Defensor EB, Pearson BL, Pobbe RLH, Bolivar VJ, Blanchard DC, Blanchard RJ. A novel social proximity test suggests patterns of social avoidance and gaze aversion-like behavior in BTBR T+ tf/J mice. Behav. Brain Res.. 2011;217:302–8.
