## Supplementary file 2 for "Oxytocin enhances acquisition in a social trust task in mice, whereas both oxytocin and its antagonist block trust violation learning"

### Results

Before conducting main analyses reported in the manuscript, or analyses on control variables reported in Supplementary file 1, we compared the male and female vehicle control (VEH) groups of the OT (PBS) and CL (DMSO) experiments (Experiments 1 and 2) on each test. If significant differences were absent, we merged the VEH groups to increase our sample size.

***Tests of explorative and anxiety-like behavior, spatial working memory and contextual learning***

**Open Field.** PBS-treated males had a significantly higher pathlength than males in the DMSO group ($w$ = 232, $p$ = .002) but spent a similar amount of time in the periphery ($t_{34}$ = 1.25, $p$ = .44) and center ($w$ = 143, $p$ = .99). PBS- and DMSO-treated females had a similar pathlength ($t_{22}$ = 0.47, $p$ = .64), and spent a similar amount of time in the periphery ($t_{22}$ = 1.76, $p$ = .18) and center ($w$ = 54, $p$ = .32). Based on these results, we merged the VEH groups in both sexes on all variables apart from male pathlength.

**T-Maze.** PBS-treated males had a significantly higher pathlength compared to DMSO-treated males over the two trials ($F_{1,22}$ = 48.75, $p$ < .001) but there were no significant novel arm preference score differences between VEH groups ($t_{22}$ = 0.88, $p$ = .39). PBS- and DMSO-treated females had a similar pathlength ($F_{1,21}$ = 0.002, $p$ = .97) and novel arm preference score ($t_{21}$ = 0.17, $p$ = .86). Based on these results, we merged the VEH groups in both sexes on all variables apart from male pathlength.

**Passive Avoidance.** Survival curves of PBS-treated males and females were not significantly different from those of DMSO-treated males ($\chi^2$ (1) = 1.45, $p$ = .23) and females ($\chi^2$ (1) = 0.68, $p$ = .41), respectively. This indicated that the proportion of mice entering the dark box (i.e., step-through latency) over time on day 2 did not differ between VEH groups for either sex. Therefore, the VEH groups were merged in both sexes.

***Tests of sociability, social memory and social safety learning***

**Social Proximity.** PBS- and DMSO-treated males engaged to the same extent in frontal contact ($t_{18}$ = 0.31, $p$ = .78) and NA behavior ($t_{18}$ = 1.11, $p$ = .78), and time spent SG ($w$ = 56, $p$ = .78). Similarly, frontal contacts ($t_{21}$ = 2.61, $p$ = .07), NA behavior ($t_{21}$ = 2.11, $p$ = .09) and time spent SG ($w$ = 60, $p$ = .92) were not different between PBS- and DMSO-treated females. Based on these results, we merged the male and female VEH groups on all variables.

**SPSN.** PBS-treated males had a significantly higher pathlength than males in the DMSO group over the three episodes of the test ($F_{1,34}$ = 6.81, $p$ = .01) but S1 ($t_{34}$ = 0.28, $p$ = .78) and S2 preference scores ($t_{34}$ = 0.89, $p$ = .38) were similar in both VEH groups. PBS- and DMSO-treated females had a similar pathlength ($F_{1,22}$ = 0.02, $p$ = .89), and S1 ($t_{22}$ = 0.11, $p$ = .92) and S2 preference scores ($t_{22}$ = 0.86, $p$ = .40). Based on these results, we merged the VEH groups in both sexes on all variables apart from male pathlength.

**STFP1.** In the *social interaction* phase, male ($t_{34}$ = 1, $p$ = .32) and female ($t_{22}$ = 0.23, $p$ = .82) VEH groups spent a similar amount of time near the demonstrator. In the *test* phase, safe food preference scores were similar in the male (time: $t_{32}$ = 0.10, $p$ = .92; food: $t_{32}$ = 0.15, $p$ = .92) and female VEH groups (time: $t_{22}$ = 0.18, $p$ = .86; food: $t_{22}$ = 0.34, $p$ = .86). Based on these results, we merged the VEH groups in both sexes on all variables.

**STFP2.** In the *social interaction* phase, male ($t_{34}$ = 0.42, $p$ = .68) and female ($t_{22}$ = 0.76, $p$ = .46) VEH groups spent a similar amount of time near the demonstrator. In the *test* phase, safe food preference scores were similar in the male (time: $t_{32}$ = 1.81, $p$ = .16; food: $t_{32}$ = 1.09, $p$ = .28) and female VEH groups (time: $t_{20}$ = 0.19 , $p$ = .85; food: $t_{20}$ = 1.58, $p$ = .26). Based on these results, we merged the VEH groups in both sexes on all variables.
